# Environmental specificity in *Drosophila*-bacteria symbiosis affects host developmental plasticity

**DOI:** 10.1101/717702

**Authors:** Robin Guilhot, Antoine Rombaut, Anne Xuéreb, Kate Howell, Simon Fellous

## Abstract

Environmentally acquired microbial symbionts could contribute to host adaptation to local conditions like vertically transmitted symbionts do. This scenario necessitates symbionts to have different effects in different environments. We investigated this idea in *Drosophila melanogaster*, a species which communities of bacterial symbionts vary greatly among environments. We isolated four bacterial strains isolated from the feces of a *D. melanogaster* laboratory strain and tested their effects in two conditions: the ancestral environment (i.e. the laboratory medium) and a new environment (i.e. fresh fruit with live yeast). All bacterial effects on larval and adult traits differed among environments, ranging from very beneficial to marginally deleterious. The joint analysis of larval development speed and adult size further shows bacteria affected developmental plasticity more than resource acquisition. This effect was largely driven by the contrasted effects of the bacteria in each environment. Our study illustrates that understanding *D. melanogaster* symbiotic interactions in the wild will necessitate working in ecologically realistic conditions. Besides, context-dependent effects of symbionts, and their influence on host developmental plasticity, shed light on how environmentally acquired symbionts may contribute to host evolution.

## Introduction

Symbiosis may contribute to host evolution through recruitment of beneficial microorganisms (Margulis and Fester 1991; Jaenike et al. 2010; Fellous et al. 2011). As the environment varies among localities, different symbionts may be most beneficial in different conditions (De Vries et al. 2004; Daskin and Alford 2012; Bresson et al. 2013; Cass et al. 2016; Couret et al. 2019), possibly explaining microbiota variation among populations of the same animal species (e.g. Chandler et al. 2011; McKenzie et al. 2017). Microbial symbionts may therefore contribute to local adaptation (Kawecki and Ebert 2004). Most studies exploring symbiont-mediated local adaptation have focused on vertically transmitted microorganisms (e.g. Moran et al. 2008). However, numerous animals form symbioses with bacteria that are in part acquired from the environment either by horizontal transmission between hosts or recruitment of free-living strains (Ebert 2013). In this context, little is known on how microbial effects on host fitness change with environmental conditions (Schwab et al. 2016; Callens et al. 2016), a necessary condition for symbiont-mediated local adaptation (Kawecki and Ebert 2004). Here, we explore how the effects of extracellular symbiotic bacteria on *Drosophila melanogaster* traits change when host and bacteria are studied in conditions that differ with their prior environment.

*Drosophila melanogaster* is a prevalent model organism for host-microbiota studies (Douglas 2018). In this species, bacterial symbionts contribute to a broad range of functions including resource acquisition, digestion, immunity and behavior (Broderick and Lemaitre 2012; Ankrah and Douglas 2018; Schretter et al. 2018). Several laboratory studies have established fly nutrition relies on interactions with gut bacteria (Shin et al. 2011; Storelli et al. 2011; Ridley et al. 2012; Wong et al. 2014; Huang et al. 2015; Leitão-Gonçalves et al. 2017; Téfit et al. 2017). In particular, bacterial genera frequently associated with laboratory flies, such as *Acetobacter* and *Lactobacillus*, can improve larval growth and development when laboratory food is poor in proteins (Shin et al. 2011; Storelli et al. 2011; Téfit et al. 2017). Even though some bacterial taxa are frequent in laboratory colonies, the composition of *Drosophila* bacterial gut communities largely varies among laboratories (Chandler et al. 2011; Staubach et al. 2013; Wong et al. 2013; Vacchini et al. 2017). Studies have shown that bacterial microbiota composition is determined by laboratory conditions more than *Drosophila* species (Chandler et al. 2011; Staubach et al. 2013), demonstrating these symbionts are largely acquired from fly environment. Empirical studies have nonetheless shown pseudo-vertical transmission of bacteria from mothers to offspring also occurs in the laboratory (Bakula 1969; Ridley et al. 2012; Wong et al. 2015; Téfit et al. 2018). Microbiota composition differences between laboratory and field flies have led authors to argue that symbiotic phenomena as observed in the laboratory may not reflect those occurring in natural conditions (Chandler et al. 2011; Winans et al. 2017). Numerous variables differ between laboratory and natural environments of *D. melanogaster* flies. A substantial difference is the composition of the nutritive substrate upon which the adults feed, copulate, oviposit and within which larvae develop. Wild flies live on and in fresh or decaying fruit flesh, usually colonized by yeast, whereas laboratory flies are reared on an artificial, jellified and homogeneous diet that contains long-chained carbohydrates (e.g. starch), agar, preservatives and dead yeast cells or yeast extract. To this date, very few studies have investigated *Drosophila*-bacteria interactions in conditions comparable to those of the field. How much *Drosophila*-bacteria interactions that occur in the laboratory are maintained in natural substrate remains largely undescribed.

Here, we experimentally studied the symbiosis between a laboratory strain of *D. melanogaster* and four bacterial symbionts (isolated from its feces) in the ancestral laboratory medium and in a new environment (grape berry) where we reproduced natural egg and bacterial deposition from mothers. After inoculating bacteria-free eggs with these four bacterial isolates, we scored various phenotypic fly traits at the larval and adult stages. We investigated two questions. (1) We focused on the influence of environmental variation on bacterial effects analyzing each of the host’s traits individually. Our aim was to unveil whether host-symbiont that occurred in the environment of origin (i.e. the laboratory) maintained in conditions more ecologically realistic. We further relate these observations to fly and bacteria ecology. (2) We performed a new, simultaneous analysis of two traits in order to disentangle symbionts’ effects on host developmental plasticity and resource acquisition, two non-excluding possibilities. Separating plasticity from resource acquisition is important for at least two reasons. First, long-term symbiotic associations would be more likely when symbionts provide new capabilities (i.e. resources) than when they affect quantitative traits (Fellous and Salvaudon 2009) or their plasticity (Chevin et al 2010). Second, recent literature shows that the evolution of symbiont transmission depends on which of host’s traits it affects (Brown and Akçay 2019); importantly, this mathematical model is based on the plastic trade-off between survival and reproduction. Recent studies have shown that in *D. melanogaster* bacteria can affect host position along this trade-off (Gould et al. 2018; Walters et al. 2018). Here, we focused on another trade-off, the relationship between duration of larval development and adult size at emergence which is well-established in holometabolous insects (Teder et al. 2014; Nunney 1996). In brief, we reasoned that bacterial effects on host developmental plasticity would move host phenotypes along the trade-off axis, while bacterial effects on resource acquisition would allow faster development or larger size without detrimental effects on the other trait (see Materials and Methods for details).

## Methods

### Drosophila strain

Insects were from the Oregon-R *Drosophila melanogaster* strain that was founded in 1927 and has since been maintained in numerous laboratories. Our sub-strain was funded ±2 years earlier from a few dozen individuals provided by colleagues. They had been reared on a laboratory medium comprising banana, sugar, dead yeast, agar and a preservative (Table S1A). Before and during the experiment reported here, all insects were maintained at 21 °C (stocks) or 23 °C (experiment), with 70 % humidity and a 14 h photoperiod.

### Microbial isolates

The starting point of this work was to isolate and cultivate symbiotic bacteria from the flies. These bacteria were chosen for their ease of cultivation and our ability to discriminate them morphologically on standard microbiological medium. Our aim was not to sample the whole community of bacteria associated with our fly stock but to carry out tractable experiments using a random subset of their symbionts. Note our isolation method excluded the *Acetobacter* spp. and *Lactobacillus* spp., some of the best known symbionts of *D. melanogaster*. However, all the bacterial strains we isolated had already been identified as associated to *Drosophila* flies (Chandler et al. 2011; Staubach et al. 2013). Available literature did point to a number of taxa which interactions with *Drosophila* flies are described, and that we could have sourced from other laboratories. However, working with strains we could readily isolate from our fly colony meant we were certain to investigate fly-bacteria associations in their environment of origin.

In order to isolate bacteria present in fly feces, several groups of twenty *Drosophila melanogaster* flies were placed in sterile glass vials for 1 h. After fly removal, vials were washed with sterile PBS (Phosphate-Buffered Saline) solution, which was then plated on Lysogeny Broth (LB) agar medium (Table S1B) and incubated at 24 °C. Four bacterial morphotypes of variable frequency were chosen based on visible and repeatable differences in size, color, general shape and transparency during repeated sub-culturing on fresh media (Figure S2). A single colony of each morphotype was amplified in liquid LB medium in aerobic conditions at 24 °C for 72 h, centrifuged and washed in PBS. Several sub-samples of equal concentration were stored at −80 °C in PBS with 15% glycerol and further used for molecular identification and the main experiment (one per experimental block).

Molecular identification of each bacterium was carried out by Sanger sequencing. To this aim, a fresh colony of each bacterial type was picked with a sterile toothpick and dipped into sterile water, then boiled 10 min at 95 °C (Mastercycler, Eppendorf) and cooled in ice water. A sterile toothpick dipped into sterile water served as sterility control of the process. Fragments of the 16sRNA gene were amplified with bacterial primers Y2MOD (5-ACTYCTACGGRAGGCAGCAGTRGG-3’) and 16SB1 (5’-TACGGYTACCTTGTTACGACTT-3’) (Haynes et al. 2003; Carletto et al. 2008). PCRs were performed in a volume of 25 µl, containing each primer at 0.2 µM, 1x buffer (containing 2 mM MgCl_2_), each dNTP at 0.2 mM, and 1 U of *DreamTaq* Taq (Thermo Scientific). PCRs cycles had an initial denaturation step at 95 °C for 15 min, followed by ten cycles at 94 °C / 40 s – 65 °C / 45 s – 72 °C / 45 s); followed by 30 cycles at 94 °C / 40 s – 55 °C / 45 s – 72 °C / 45 s; and finished with an extension step of 10 min at 72 °C. Negative PCR controls were included. PCR products were visualized under UV light in an agarose gel before sequencing. Consensus sequences were created with CodonCode Aligner 4.2.7. Online SINA alignment service (https://www.arb-silva.de/aligner/) (Pruesse et al. 2012) and NCBI GenBank blastn service (https://blast.ncbi.nlm.nih.gov/Blast.cgi) were used to compare and assign the sequences. The four bacteria were identified as a *Staphylococcus* (likely *S. xylosus*), an *Enterococcus* (likely *E. faecalis*), an Enterobacteriaceae and an Actinobacteria (likely *Brevibacterium*). Further in this article, theses bacteria are referred to as *Staphylococcus*, *Enterococcus*, Enterobacteriaceae and Actinobacteria, respectively. All sequences were deposited in the NCBI database under the accession numbers MK461976 (*Staphylococcus*), MK461977 (*Enterococcus*), MK461978 (Enterobacteriaceae) and MK461979 (Actinobacteria).

A wild isolate of *Saccharomyces cerevisiae* yeast was used in experiments where larvae developed in fresh grape berries. The yeast was isolated from a wild Drosophilid in a vineyard in Southern France (*‘Le Domaine de l’Hortus’*, Hérault, France) (see Hoang et al. (2015) for a balanced discussion on *Drosophila-Saccharomyces* interactions). The isolate was grown in YPD medium, washed, split into several samples, stored at −80 °C in sterile PBS with 15 % glycerol, that were further used in the experiment (one per block).

### Experimental design

Flies were associated with bacteria following a full-factorial design that resulted in twelve different treatments. There were two types of fly environments: laboratory medium (the ancestral environment, see Table S1A for composition) and grape berries (the new environment, white grapes, unknown cultivar). We had six different symbiont treatments: each of the four bacterial strains described above, a mix of the four bacteria and controls without bacteria (Figure 1). Each treatment had 13 to 15 replicates organized in 15 blocks launched over four days. Bacterial growth was also studied in fly-free grapes but is not described here.

**Figure 1:**
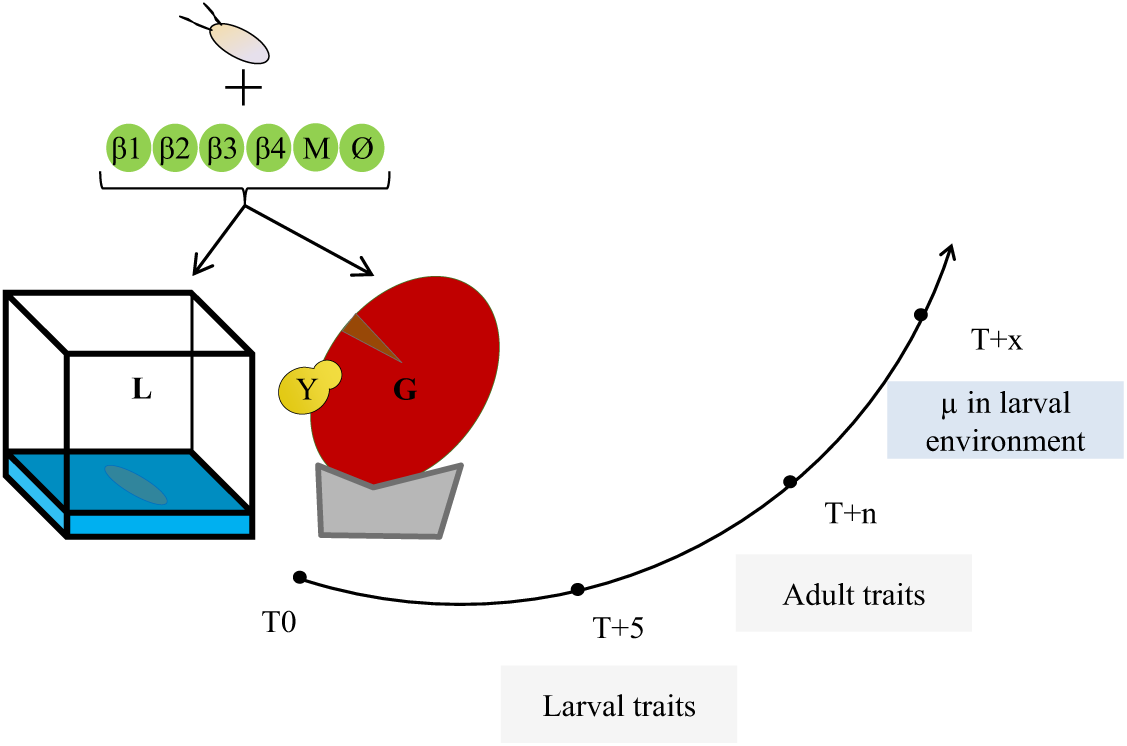
summary of the experimental design and the measured traits. T0: association of Drosophila eggs with bacteria (β1, β2, β3, β4), bacterial mixture (M), or nothing (Ø), in the two environments: laboratory medium (L) or grape berry (G) inoculated with live yeast (Y). T+5: larval traits scoring after five days. T+n: adult size scoring on a randomly chosen subset of adults from each replicate. T+x: analysis of the microbial content of the larval environment two days after the end of pupal formation.

Grape berries were surface-sterilized in a 2 % bleach solution before use. Because *D. melanogaster* females only oviposit in wounded fruit, we incised 5 mm of berry skin (Figure S4) where we deposited twenty eggs free from culturable bacteria. These eggs were produced by the oviposition of flies on laboratory medium supplemented with the antibiotic streptomycin (1 mg / ml in 1 mM EDTA, Sigma-Aldrich ref. 85886). The efficacy of this method for removing culturable bacteria from egg surface was confirmed by the lack of bacterial growth after the deposition of such eggs onto LB agar plates (note however that these conditions were not suitable for detection of anaerobic bacteria such as *Lactobacillus*). Grape berries were inoculated with live yeast cells as it is a key nutritional component (Begg and Robertson 1948; Becher et al. 2012) and was necessary for fly survival in our system (Figure S3). For treatments with laboratory diet we deposited twenty eggs free from culturable bacteria on incisions at the surface of 4 ml of medium placed in 2 cm * 2 cm plastic cubes. Berries and laboratory media were all placed in 75 ml plastic vials closed by a foam plug.

Bacterial cells were inoculated to laboratory medium and grape berry immediately before egg deposition. Single bacterial strain treatments received 10^4^ live bacterial cells, and the mixed treatment 2.5 × 10^3^ cells of each bacterium (i.e. 10^4^ cells in total), suspended in 10 µl of sterile PBS. The number of inoculated bacterial cells was chosen based on the average number of bacteria previously reported in the guts of second-instar *Drosophila* larvae (Bakula 1969; Storelli et al. 2011). In control treatments, sterile PBS was deposited instead of bacteria. On grape berries, 10^4^ live cells of the yeast *Saccharomyces cerevisiae* were inoculated. Note fruit substrate and live yeast presence are confounded factors in our experiment because we did not intend to study the effect of live yeast onto larval growth (Becher et al 2012) but to simulate field conditions where larvae develop in presence of live yeast. Although the laboratory medium also contains yeast (Table S1A), cells are killed during industrial production.

### Fly phenotyping

We scored six different phenotypic traits in larvae and adults: larval size after five days; larval mouthpart movement rate after five days; visible number of larvae on medium surface after five days; survival rate to adult emergence; time until adult emergence and a proxy of adult size. Larval mouthpart movement speed was the number of back-and-forth movements of the mouthpart that could be observed in five seconds. Newly formed pupae were transferred to empty sterile vials daily. We recorded male and female emergences daily.

The size of adults, and their microbial content (see below), were estimated on a subset of adults that emerged from each vial. For each vial, one pupa was chosen randomly and all adults that emerged on the same day as the focal pupa were collected and pooled by sex. These pools were homogenized in 200 µl of sterile PBS using a sterile pestle, splat in two sub-samples and stored at −80 °C with 15 % sterile glycerol. One of the two sub-samples was used to numerate live bacteria and yeast cells in newly emerged adults. The other sub-sample was used to estimate adult size with the spectrophotometric method described in Fellous et al. (2018). We chose this method as it allowed the simultaneous analysis of adult size and microbial content. Briefly, we used log-transformed optical density at 202 nm of fly homogenate as a proxy of adult size. Optical density of homogenates was measured several months after the experiment when samples were thawed, crushed a second time using a Tissue Lyser II (Qiagen) for 30 s at 30 Hz with Ø3 mm glass balls, centrifuged for 30 s at 2000 G. Optical density of 15 µL of supernatant was then read on a Multiskan GO spectrometer (Thermo Scientific). This metrics correlates in both males and females with wet weight and wing length (all R^2^ > 0.8), two frequently used size proxies in *Drosophila* studies. For figures and analyses of adult size we used the Log_10_ of observed optical density divided by the number of individuals in the sample.

### Analysis of bacterial presence and metabolism

We tested the presence of inoculated bacteria and yeast in substrates two days after the appearance of the last pupa. Samples were analyzed by plating homogenates on LB agar medium and incubated at 24 °C. In this manuscript we only report on the presence or absence of inoculated bacteria in the larval substrate. Data of microorganism presence and numbers in emerging adults will be reported separately. The Enterobacteriaceae and the Actinobacteria were the main bacterial strains that affected fly phenotypes. In order to shed light on the ecologies of these two strains and therefore on their effects on hosts, we analyzed their metabolic capabilities with Eco Microplates (Biolog) (see Text S5 for methodological details).

Bacteria and fungi morphologically different from those we had inoculated were observed in samples from 17 % of the vials (either in adults or in the environment). Data from these vials were excluded for all analyzes presented here. Both datasets are available in the open data repository Zenodo (DOI: 10.5281/zenodo.2554194).

### Statistical analyses

#### Individual traits

To study the response of each fly phenotypic trait to variation of larval substrate and bacterial symbiont, we used linear mixed models (LMM) with Restricted Maximum Estimate Likelihood (REML). Fixed factors were the ‘larval environment’ (i.e. laboratory medium or fruit), ‘bacterial treatment’, ‘fly sex’ (for the analyses of age at emergence and adult size only), and their full-factorial interactions. ‘Block identity’ was defined as random factor in all models and a random term indicating the vial in which the flies developed was added to the analysis of age at emergence. A Backward, stepwise model selection was used to eliminate non-significant terms from initial, full models. Homoscedasticity and residuals normality visually complied with model assumptions. When the ‘bacteria*environment’ interaction was significant, and to investigate hypotheses based on the visual observation of the data, we used independent contrasts to test significant differences between bacterial treatments and controls from the same environment.

#### Joint effect of bacteria on adult age and size at emergence

The aim of this analysis was to study how bacteria affected simultaneously speed of larval development and adult size. Importantly, we needed to discard the general effect of the nutritive environment to single out the effects of the symbionts. Indeed, if one environment was generally more favorable than the other, main environmental effects could create a positive relationship between the two traits that would conceal how bacteria affect simultaneously the two traits. To this end, all analyses were carried out after subtracting the mean trait value of the controls (i.e. bacteria-free) in the relevant environment from the trait values of each combination of bacteria and environment. In other words, data presented in Figures 5 and S6 represent the incremental effects of the bacteria on host traits after removal of the overall influence of the nutritive substrate.

We carried out two types of analyses. (1) In order to unveil the overall pattern (Figures 5 and S6) we worked with mean treatment effects (i.e. one single data point per treatment, two when sex was taken into account) and univariate regressions. Because of the significant interaction between sex, bacteria and environment for adult size, our initial analysis separated males from females (Figure S6). However, the linear regression of size onto developmental speed was not significantly different among sexes (Interaction Sex*Speed: F_1,16_ = 2.93, p = 0.11). Presented results hence merge observations from males and females. (2) In order to explain the factors behind the simultaneous effect of bacteria on developmental speed and adult size we carried out a multivariate analysis of variance (MANOVA) using all data points (i.e. one data point per experimental unit). MANOVA was chosen because it enables studying how factors affect several variables jointly, in other words it considers effects onto the correlation between several variables (Zar 2009, p.319). We used a “repeated measures” personality of MANOVA and reported the tests based on the Sum response function (i.e. a M-matrix that is a single vector of 1 s; between-subject report in JMP). Model contained the factors ‘bacterial treatment’, ‘environment’ and their interaction. Homoscedasticity and residuals normality visually complied with MANOVA assumptions. The dataset used for the MANOVA analysis is available in the open data repository Zenodo (DOI: 10.5281/zenodo.3352230).

Analyzes were performed with JMP (SAS, 14.1).

## Results

### Effects of bacteria on individual traits reveal extensive environmental-dependence

*Larval size* after five days was influenced by an interaction between the environment and the bacterial treatment (Table 1, Figure 2A). In grapes, addition of the Actinobacteria decreased larval size relative to bacteria-free controls but had no effect in laboratory media. In laboratory media, addition of the Enterobacteriaceae alone or in mixture with the other bacterial strains produced larger larvae than bacteria-free controls (contrast ‘Enterobacteriaceae treatment’ vs ‘Control treatment’: F_1,90_ = 28.92, p < 0.0001), which did not happen when grown on a grape substrate (contrast ‘Enterobacteriaceae treatment’ vs ‘Control treatment’: F_1,86_ = 0.92, p = 0.3405) (Figure 2A).

**Figure 2:**
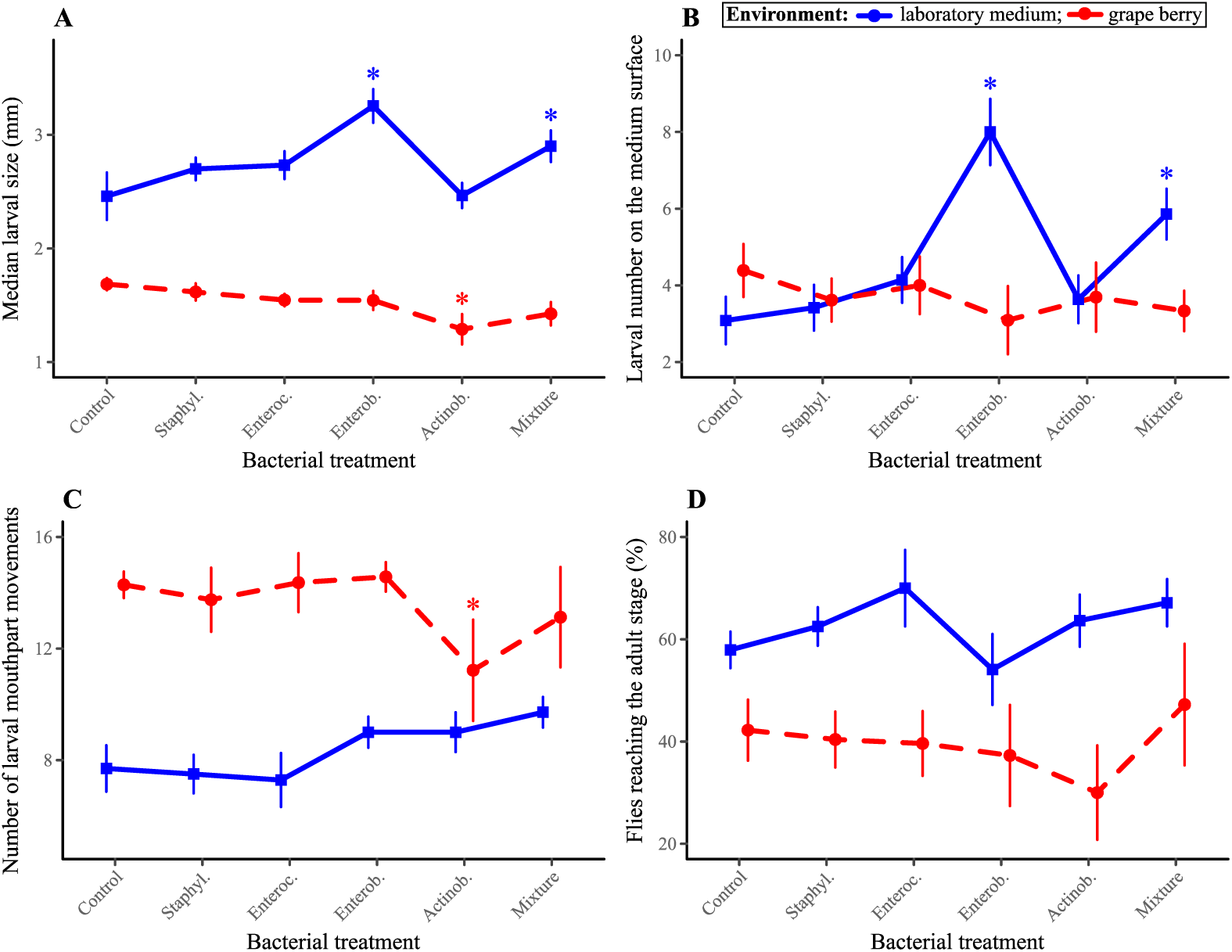
larval phenotypes in response to bacterial treatment and larval environment. (A) Median larval size after five days; (B) Number of larvae on the medium surface after five days; (C) Number of larval mouthparts movements per five seconds observed after five days; (D) Survival from egg to adult. Symbols indicate means; error bars indicate standard errors around the mean. Stars (*) indicate treatments significantly different from controls in the same environment (post-hoc contrasts, α = 0.05).

**Table 1:**
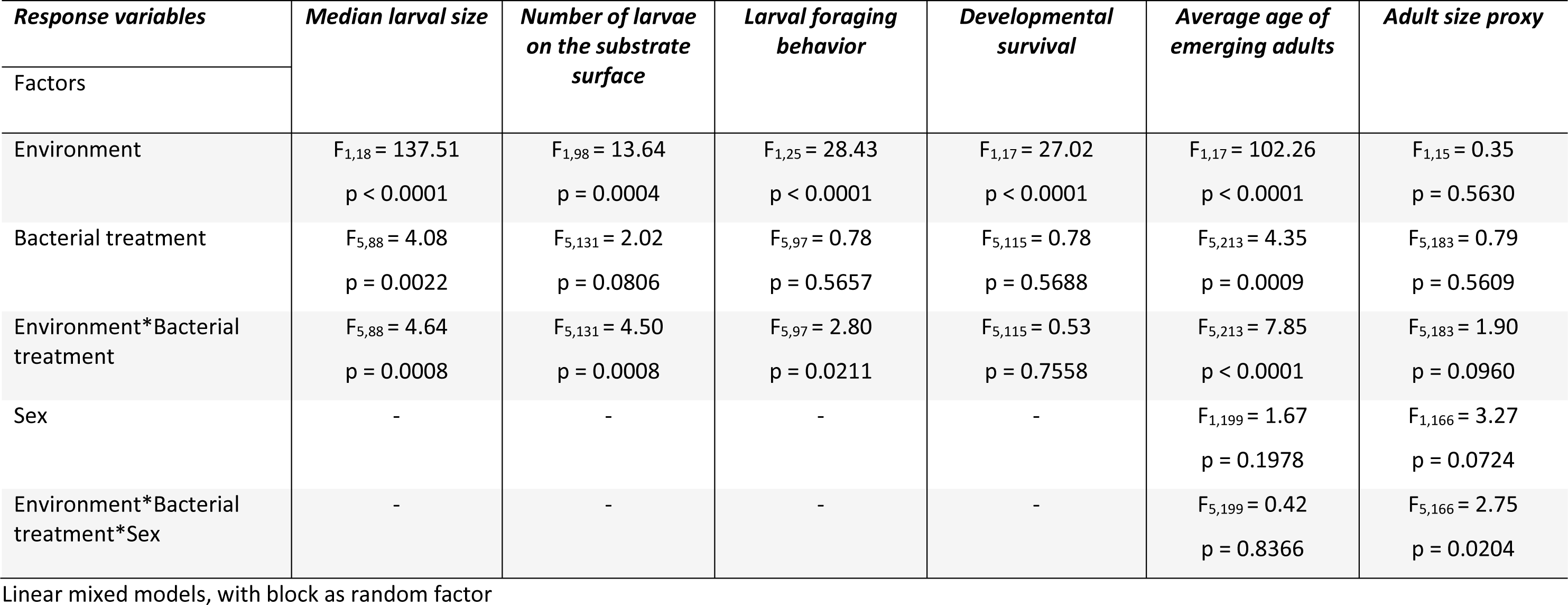
analysis of larval and adult phenotypes in response to bacterial treatment and larval environment. Linear mixed models (REML).

| *Response variables* | *Median larval size* | *Number of larvae on the substrate surface* | *Larval foraging behavior* | *Developmental survival* | *Average age of emerging adults* | *Adult size proxy* |
| --- | --- | --- | --- | --- | --- | --- |
| Factors |  |  |  |  |  |  |
| Environment | $F_{1,18} = 137.51$<br>$p < 0.0001$ | $F_{1,98} = 13.64$<br>$p = 0.0004$ | $F_{1,25} = 28.43$<br>$p < 0.0001$ | $F_{1,17} = 27.02$<br>$p < 0.0001$ | $F_{1,17} = 102.26$<br>$p < 0.0001$ | $F_{1,15} = 0.35$<br>$p = 0.5630$ |
| Bacterial treatment | $F_{5,88} = 4.08$<br>$p = 0.0022$ | $F_{5,131} = 2.02$<br>$p = 0.0806$ | $F_{5,97} = 0.78$<br>$p = 0.5657$ | $F_{5,115} = 0.78$<br>$p = 0.5688$ | $F_{5,213} = 4.35$<br>$p = 0.0009$ | $F_{5,183} = 0.79$<br>$p = 0.5609$ |
| Environment*Bacterial treatment | $F_{5,88} = 4.64$<br>$p = 0.0008$ | $F_{5,131} = 4.50$<br>$p = 0.0008$ | $F_{5,97} = 2.80$<br>$p = 0.0211$ | $F_{5,115} = 0.53$<br>$p = 0.7558$ | $F_{5,213} = 7.85$<br>$p < 0.0001$ | $F_{5,183} = 1.90$<br>$p = 0.0960$ |
| Sex | - | - | - | - | $F_{1,199} = 1.67$<br>$p = 0.1978$ | $F_{1,166} = 3.27$<br>$p = 0.0724$ |
| Environment*Bacterial treatment*Sex | - | - | - | - | $F_{5,199} = 0.42$<br>$p = 0.8366$ | $F_{5,166} = 2.75$<br>$p = 0.0204$ |

Linear mixed models, with block as random factor

*The number of larvae visible on medium surface* was influenced by an interaction between the environment and the bacterial treatment (Table 1, Figure 2B). In laboratory media, addition of the Enterobacteriaceae alone or in mixture with the other bacterial strains led to greater numbers of visible larvae compared to bacteria-free controls (contrast ‘Enterobacteriaceae treatment’ vs ‘Control treatment’: F_1,131_ = 20.40, p < 0.0001; contrast ‘Mixture treatment’ vs ‘Control treatment’: F_1,131_ = 6.98, p = 0.0092), which did not happen when grown on a grape substrate (contrast ‘Enterobacteriaceae treatment’ vs ‘Control treatment’: F_1,131_ = 1.63, p = 0.2036; contrast ‘Mixture treatment’ vs ‘Control treatment’: F_1,131_ = 0.93, p = 0.3355) (Figure 2B).

*Mouthparts movement rate* was influenced by an interaction between the environment and the bacterial treatment (Table 1, Figure 2C). Movements were generally faster in grapes than in laboratory media. However, addition of the Actinobacteria slowed down the movements of mouthparts in grapes to a level comparable to the one of larvae reared on laboratory media (contrast ‘Actinobacteria treatment’ vs ‘Control treatment’: F_1,99_ = 4.54, p = 0.0355) (Figure 2C).

*The proportion of eggs surviving until the adult stage* was only affected by the environment, with a lower survival in grapes than in laboratory media (Table 1, Figure 2D). Even in laboratory medium, where survival was best, it never exceeded 70%. We believe a fraction of the eggs were hurt during experiment set-up.

*Age at adult emergence* was not different among sexes but was influenced by an interaction between the environment and the bacterial treatment (Table 1, Figure 3). In laboratory media, flies reared with the Enterobacteriaceae, alone or in mixture, emerged nearly two days sooner than bacteria-free flies in the same environment and almost four days earlier than bacteria-free flies in grapes (contrast ‘Enterobacteriaceae treatment’ vs ‘Control treatment’: F_1,229_ = 27.20, p < 0.0001; contrast ‘Mixture treatment’ vs ‘Control treatment’: F_1,227_ = 24.36, p < 0.0001) (Figure 3). In grapes, flies reared with the bacterial mixture emerged one day later than bacteria-free flies (contrast ‘Mixture treatment’ vs ‘Control treatment’: F_1,226_ = 6.21, p = 0.0135) (Figure 3).

**Figure 3:**
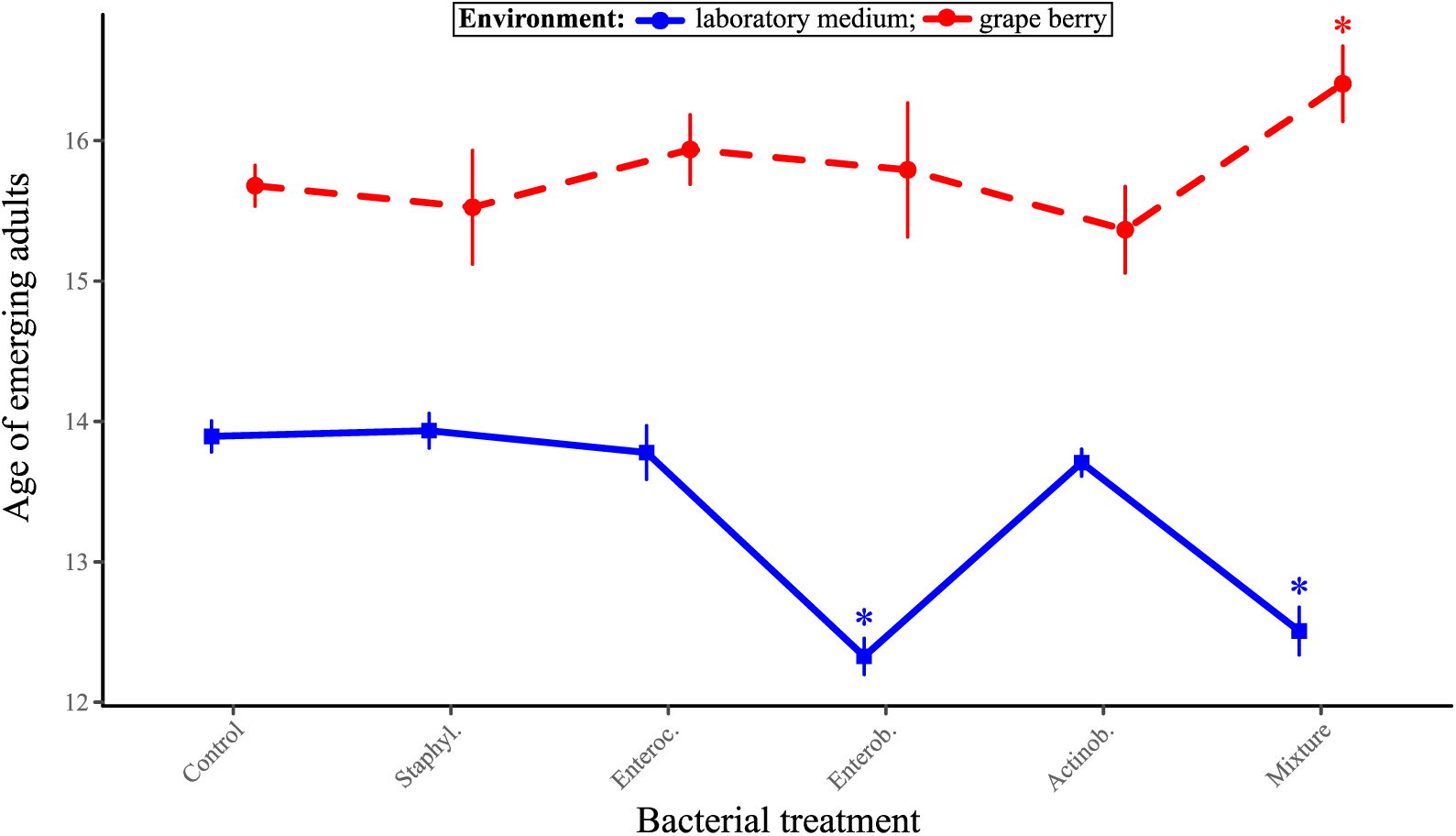
age of *Drosophila* adults at emergence in response to bacterial treatment and larval environment. Symbols indicate means; error bars indicate standard errors around the mean. Stars (*) indicate treatments significantly different from controls in the same environment (post-hoc contrasts, α = 0.05).

*Adult size* was influenced by the triple interaction between sex, the environment and the bacterial treatment (Table 1, Figure 4). Several bacterial treatments had sex-specific effects that differed among the two environments. For example, inoculation of the mixture of the four bacteria produced larger males than females in grapes (contrast ‘Mixture treatment’ vs ‘Control treatment’: F_1,166_ = 5.30, p = 0.0225), but smaller males than females in laboratory media (contrast ‘Mixture treatment’ vs ‘Control treatment’: F_1,167_ = 4.79, p = 0.0300) (Figure 4). Similarly, inoculation of the *Staphylococcus* or *Enterococcus* led to larger males than females in grape (contrast ‘*Staphylococcus* treatment’ vs ‘Control treatment’: F_1,164_ = 4.97, p = 0.0271; contrast ‘*Enterococcus* treatment’ vs ‘Control treatment’: F_1,164_ = 7.48, p = 0.0069), but no difference in laboratory medium (contrast ‘*Staphylococcus* treatment’ vs ‘Control treatment’: F_1,165_ = 0.11, p = 0.7367; contrast ‘*Enterococcus* treatment’ vs ‘Control treatment’: F_1,167_ = 0.66, p = 0.4182) (Figure 4).

**Figure 4:**
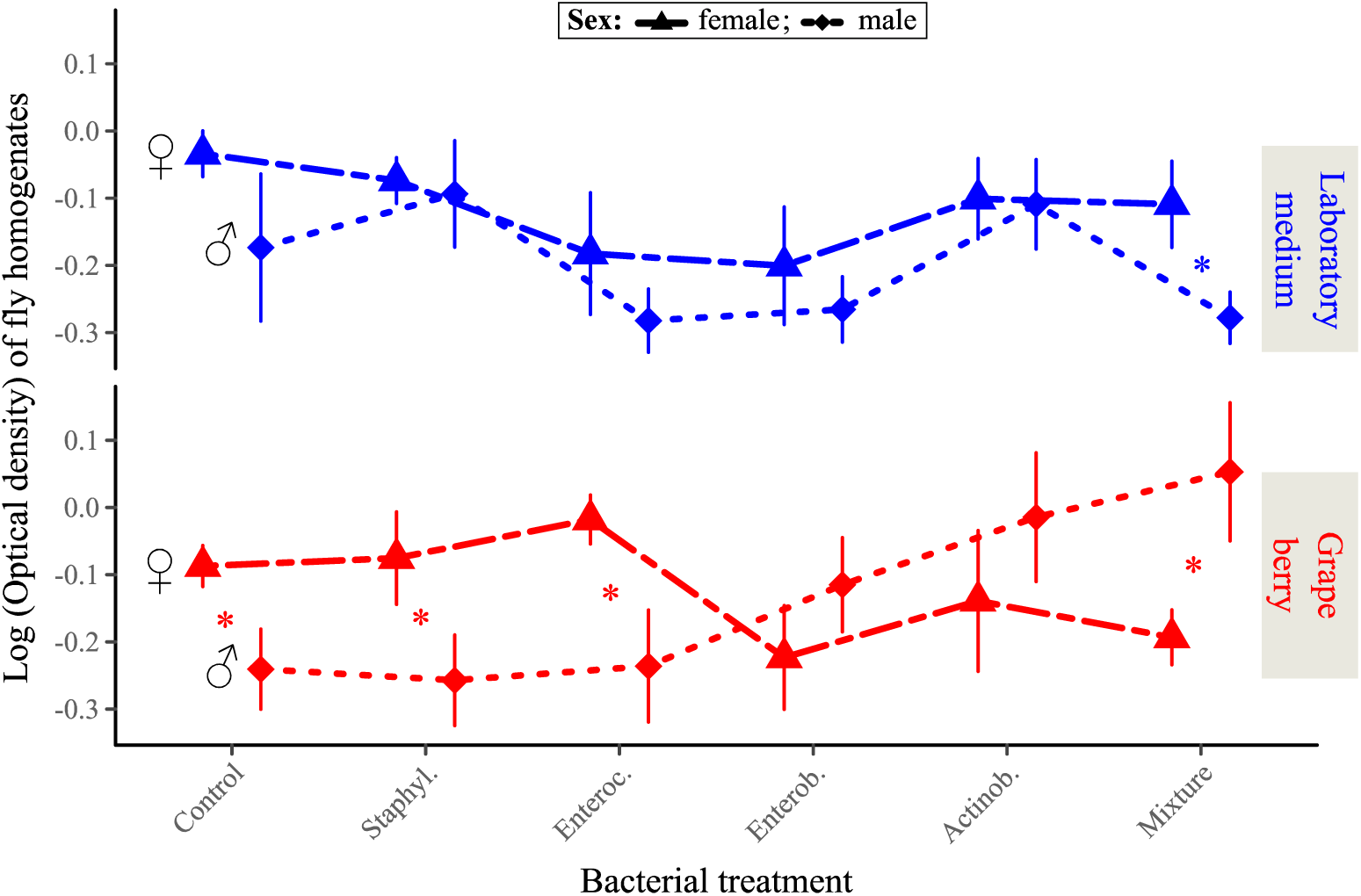
***Drosophila* adult size proxy in response to bacterial treatment and larval environment.** Symbols indicate means; error bars indicate standard errors around the mean. Stars (*) indicate significant differences between males and females in the same environment (post-hoc contrasts, α = 0.05).

**Figure 5:**
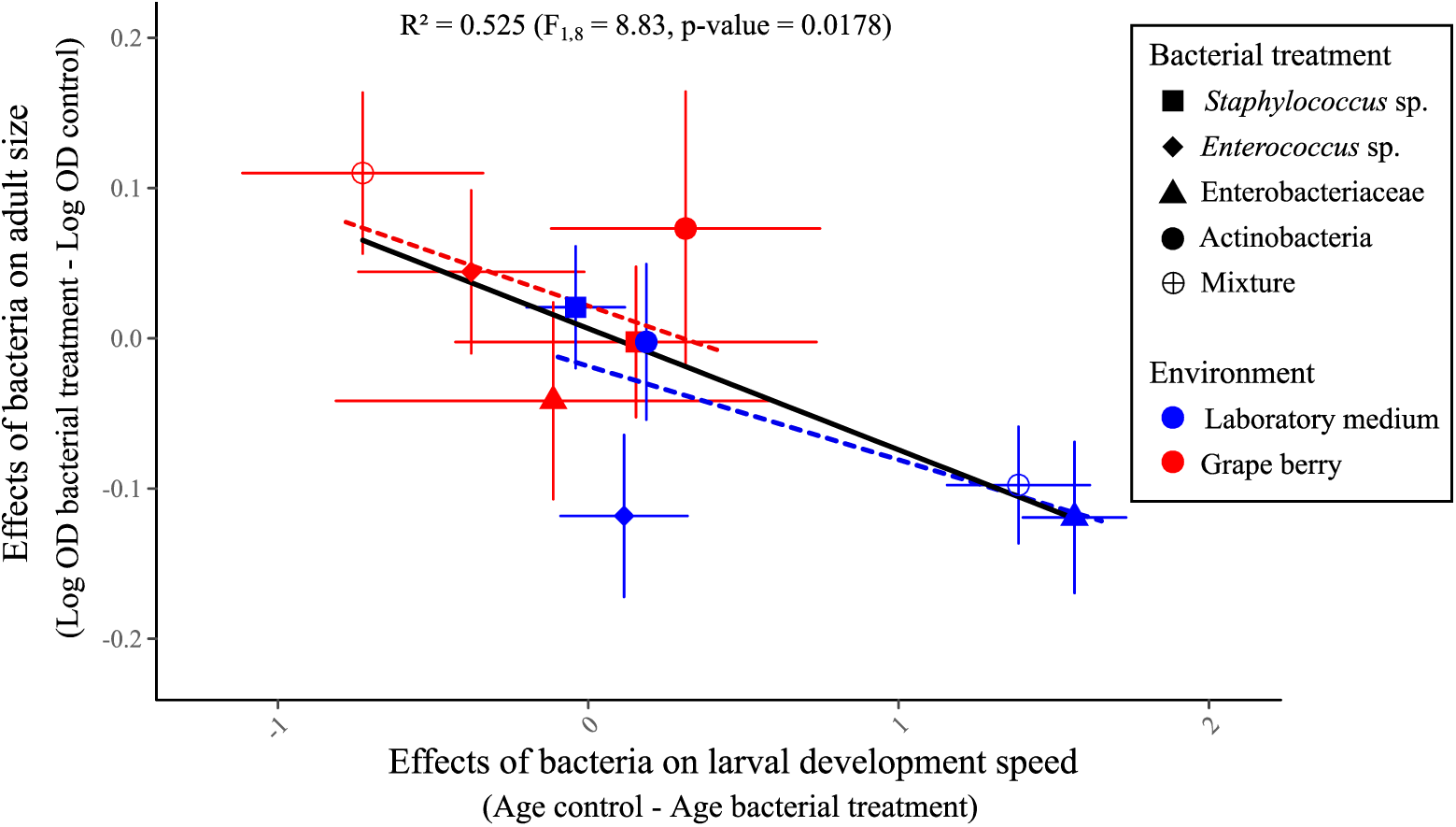
relationship between bacterial effects on age of emerging adults and bacterial effects on adult size. Effects of bacteria for each treatment were calculated by subtracting the mean trait value of controls in the same environment to mean trait value of the treatment. Error bars indicate standard errors around the means. The dashed regression lines represent the relationships between the two traits in each environment.

**Figure 6:**
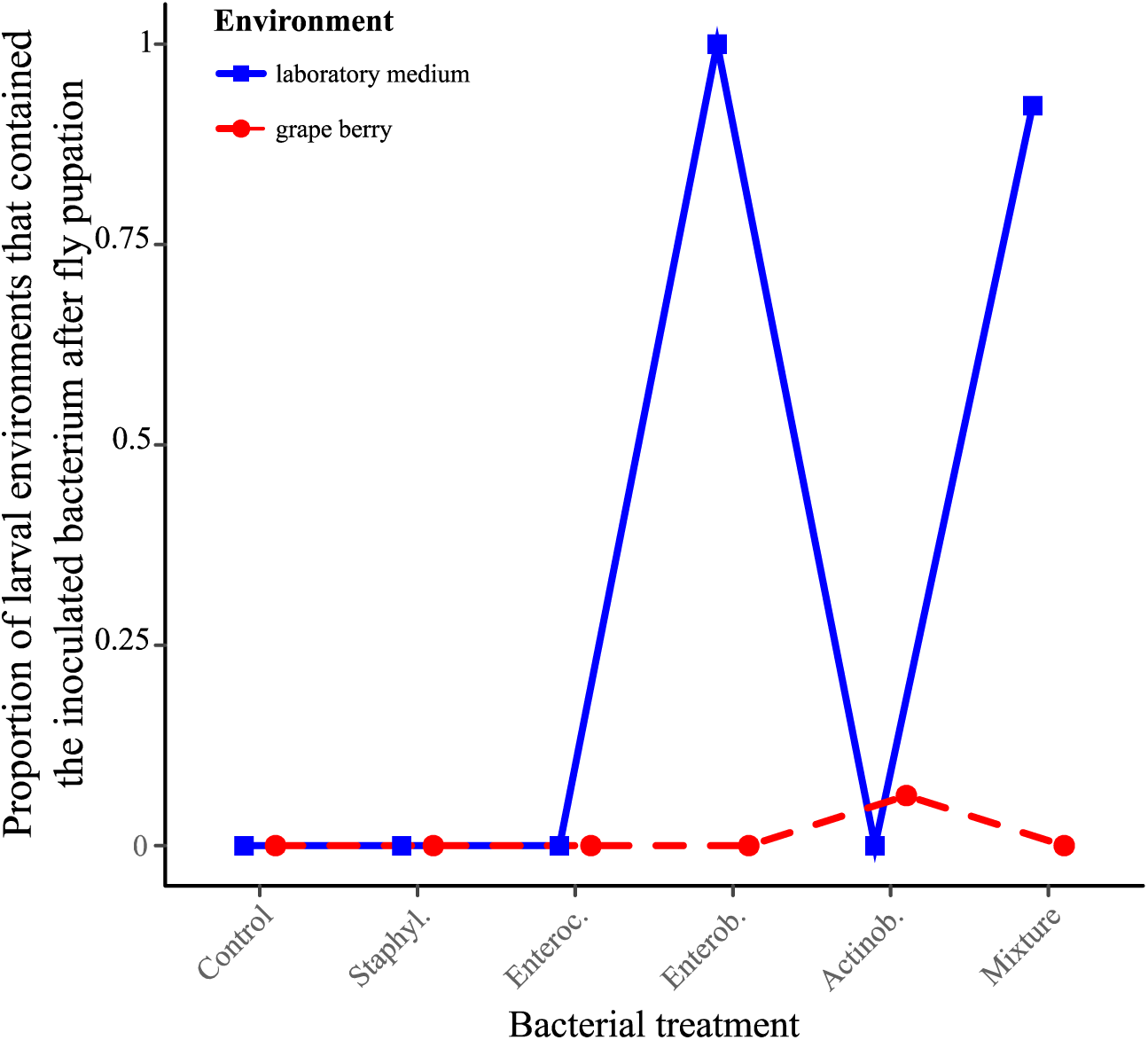
proportion of larval environments that contained the inoculated bacterium two days after the formation of the last pupa. Proportions were calculated over 7-26 replicates: Lab. (Laboratory medium) – Control (n = 12 replicates), Lab. – Staphyl. (n = 11), Lab. – Enteroc. (n = 7), Lab. – Enterob. (n = 10), Lab. – Actinob. (n = 10), Lab. – Mixture (n = 13), Grape – Control (n = 26), Grape – Staphyl. (n = 16), Grape – Enteroc. (n = 16), Grape – Enterob. (n = 13), Grape – Actinob. (n = 16), Grape – Mixture (n = 12).

### Joint analysis of adult age and size at emergence suggests bacteria affect host developmental plasticity along a trade-off

We expected three possible patterns when plotting average adult size in function of speed of larval development (i.e. – age at emergence): a positive correlation indicative of a similar effect of the bacteria on the two traits (i.e. bacteria mostly modulate resource acquisition); a negative correlation indicative of bacteria affecting host position along the trade-off (i.e. bacteria mostly modulate developmental plasticity); a lack of correlation that would have been challenging to interpret on its own as several processes could produce this result (e.g. bacterial effects on both host plasticity and resource acquisition).

The relationship between effects of bacteria on adult age and size at emergence was marginally significant and negative (Linear model F_1,8_ = 8.83, p = 0.018) (Figure 5). A MANOVA shed light on the relative influence of the environment and the bacterial treatments on the correlated effect of the treatments on the two traits (Table 2) (see Table S6 for MANOVA results for males and females). It revealed the environment was an important factor: in laboratory medium, addition of bacteria accelerated development relative to controls at the cost of producing smaller adults; in grape addition of bacteria slowed down development relative to controls but emerging adults were large (Figure 5). There was no significant main effect of the bacterial treatments but a significant interaction with the environment, which confirms the bacterial treatments had different effects on host phenotype in each environment. Analyzing the relationships between the two traits with MANOVA in each environment separately (Figure 5, dashed regression lines) revealed a significant effect of the bacteria in laboratory medium (F_4,46_ = 13.9, p <0.0001) but not in grape (F_4,39_ = 0.55, p= 0.7).

**Table 2:** Multivariate Analysis of Variance of the joint effect of the bacteria on ‘Age at emergence’ and ‘Adult size’. As in Figure 5, general effects of the environments were removed by subtracting trait values of controls (i.e. without bacterial addition) in each environment before carrying out the analysis.

| *Factor* | *F* | *d.f.* | *p* |
| --- | --- | --- | --- |
| Environment | 14.9 | 1.85 | 0.0002 |
| Bacterial treatment | 1.65 | 4.85 | 0.17 |
| Environment*Bacterial treatment | 3.86 | 4.85 | 0.006 |

### Bacterial occurrence in the environment and their metabolism

The Enterobacteriaceae isolate was the only bacterium to be consistently retrieved from the laboratory medium in which larvae had developed (Figure 6). In one instance, the Actinobacteria was found in a grape berry from which no live adult fly emerged (Figure 6). The physiological profile of the Enterobacteriaceae revealed growth of the bacterium in a broad panel of carbon sources (Figure S5A). The physiological profile of the Actinobacteria revealed substantial growth of the bacterium on the carbon sources Pyruvic Acid Methyl Ester and Tween 80 only (Figure S5B).

## Discussion

We studied the symbiotic interactions between a laboratory strain of *Drosophila melanogaster* and four bacterial strains isolated from its feces. Our results show different effects of bacterial symbionts on host phenotype in laboratory medium and in real fruit. All symbiont effects were environment-dependent, some of which may be explained by the ecology of laboratory-associated symbionts in artificial medium. The joint analysis of larval developmental speed and adult size further suggests bacteria influence host developmental plasticity along the well-known physiological trade-off between the two traits.

### Different symbiont effects in different environments

The observation that all bacterial effects on host phenotype were different in laboratory medium and grape berry prompts the question of the reason behind this discrepancy. Focusing of the Enterobacteriaceae may shed light onto the ecologies of the symbiotic bacteria we isolated, and why they differed among environments.

In laboratory medium, inoculation of the Enterobacteriaceae induced greater larval size and accelerated larval development (Figures 2A and 3). Besides, adults produced by larvae associated with the Enterobacteriaceae in laboratory medium were not significantly smaller than adults produced by bacteria-free larvae (Figure 4). The bacterium hence accelerated larval growth. In its presence larvae could be observed in greater numbers at the surface of the medium than in the absence of the bacterium (Figure 2B), even though there were no mortality differences among Enterobacteriaceae-associated and bacteria-free larvae (Figure 2D). The Enterobacteriaceae was also the only bacterium to be retrieved from the medium after fly pupation (Figure 6). These elements may be explained by three mechanisms. (1) The numerous larvae observed on laboratory medium surface in presence of the Enterobacteriaceae could be a direct consequence of accelerated development. Indeed, larvae at the end of the third instar are often referred to as ‘wandering larvae’ because they move out of the larval environment in search of a place to pupate. (2) The bacterium could serve as food and be grazed on medium surface by foraging larvae. The phenomenon would be similar to that described by Yamada et al. (2015) where the yeast *Issatchenkia orientalis* extracts amino acids from agar-based laboratory medium and concentrates them on medium surface where adult flies harvest them. This hypothesis is congruent with the visual observation that media inoculated with the Enterobacteriaceae harbored white microbial growth on their surface (Figure S7). Along these lines, the wide metabolic spectrum of this bacterium (Figure S5A) suggests the microorganism is a generalist that would extract resources from the medium, possibly transform nutrients (Ankrah and Douglas 2018; Sannino et al. 2018), and eventually concentrate them on medium surface. (3) Microbial growth at the surface would interfere with larval development in such way that larvae would remain at the surface. This behavior could also trigger accelerated development if excessive microbial growth revealed detrimental. The three hypotheses above are non-excluding; the joint-analysis of developmental speed and adult size sheds further light on this question (see below).

Why did the effect of the Enterobacteriaceae on host phenotype differ among environments? The physical nature of laboratory medium is very different from that of real fruit. In particular, the agar of laboratory medium permits the diffusion of simple nutrients and their absorption by bacteria and yeast present on surface. Besides, in grape nutrients are not free to diffuse but enclosed in cells. Surface growth is therefore more likely in artificial medium than in grape berry, leading to different effects on larval development. In addition to physical differences between laboratory medium and fresh fruit, the nature and concentration of available nutrient are likely to differ. It is well known that lactic and acetic acid bacteria, two taxa that were not investigated in our experiment, can promote larval growth upon nutrient scarcity (Shin et al. 2011; Storelli et al. 2011, Téfit et al. 2017). However, it is also well established that bacteria can affect *Drosophila* phenotype through signaling (Storelli et al. 2011) as well as nutrient provisioning (Brownlie et al. 2009; Bing et al. 2018; Sannino et al. 2018). In most cases, these effects which were described from laboratory flies and in laboratory medium, are condition specific (Douglas 2018). Indeed, bacteria are often only beneficial when laboratory food has a low concentration in dead yeast (i.e. amino acids) (Shin et al. 2011; Storelli et al. 2011). Along these lines, it may seem paradoxical the Enterobacteriaceae only accelerates larval growth in rich laboratory medium rather than in grape berry (unless the bacterium synthesized a rare nutrient). Metabolic profiling (Figure S5A) further shows the Enterobacteriaceae is a generalist bacterium able to grow on a variety of substrate. However, the Actinobacteria had a narrower metabolic spectrum (Figure S5B), suggesting it is a specialist which growth largely depends on the availability of specific nutrients. The bacterium slowed down larval growth in grape (Figure 2A) for an unknown reason – maybe because it exerted additional stress onto larvae in a relatively poor medium – but had no notable effect in laboratory medium. The environment-specific effect of the Actinobacteria compares to previous reports of *Drosophila* symbionts being beneficial in some environments only (e.g. *Lactobacillus plantarum* in rich medium), and further reveals that bacteria with little effect in an environment can become detrimental in new conditions.

### Effects of bacteria on host developmental plasticity

In holometabolous insects, the duration of larval development and adult size are often negatively correlated due to a physiological trade-off: faster development reduces the duration of food intake and leads to smaller adult size (Teder et al. 2014; Nunney 1996). We propose to exploit this trade-off to separate symbionts’ effects on host developmental plasticity and resource acquisition. As discussed above, symbionts of *Drosophila* flies can modify host’s signaling (e.g. Shin et al. 2011; Storelli et al. 2011), modify the nature of the larval environment as well as provide rare resources directly to the host (e.g. Brownlie et al. 2009; Sannino et al. 2018) or through the substrate. These mechanisms are expected to have different effects on the trade-off between speed of development and size. For example, effects of bacteria on signaling would move hosts along the trade-off, while the provisioning of greater resources should enable faster growth and/or larger size without sacrificing the other trait. To investigate symbionts’ effects on host developmental plasticity and resource acquisition, we extracted bacterial effects on host phenotype by subtracting control trait values to those of each of the bacterial treatments in each environment. The resulting plot of symbionts effects on developmental speed and adult size (Figures 5 and S6) reveals the influence of the bacteria on the host independently of the general effects of the environment (i.e. those not due to the bacteria).

Our original analysis of bacterial effects on larval development and adult size revealed a negative relationship (Figures 5 and S6). Treatments that accelerated development produced small adults and treatments that slowed down development produced large adults. Results suggest bacterial treatments influenced host development plastically along the trade-off between speed of development and adult size. This observation echoes recent findings showing that *Drosophila* bacterial symbionts may induce a trade-off between lifespan and fecundity (Gould et al. 2018; Walters et al. 2018). On the other hand, our results contrast with previous reports on *Drosophila* bacterial and yeast symbionts that induce positive relationships between larval and adult traits (Anagnostou et al. 2010; Bing et al. 2018; Pais et al. 2018). For example, some bacterial symbionts can positively affect both speed of larval development and adult fecundity (Pais et al. 2018). Furthermore, the yeast *Metschnikowia pulcherrima* produces small adults that are also slow to develop (Anagnostou et al. 2010). Different symbionts in different contexts can therefore affect host developmental plasticity or host resource acquisition.

The visual examination of Figure 5 shows bacterial effects measured in laboratory medium (blue points) group in the fast development-small size region of phenotypic space, while effects in grape (red points) occur in the small speed-large size side of the trade-off. This suggests that the environment could determine whether bacteria accelerate development (at the cost of a smaller size) or favor size (at the cost of a slower development). A MANOVA revealed a significant effect of the environment on the joint analysis of the two traits, hence confirming that bacterial influence on host developmental plasticity is largely determined by the environment. With only five bacterial treatments per environment it was not possible to test if bacteria affect host development along the trade-off within a single environment.

Whether microbial symbionts influence hosts through effects on developmental plasticity or resource availability (i.e. general vigor *sensus* Fry (1993)) may change the evolutionary fate of the host-symbiont relationship. First, symbionts that plastically alter phenotypes would be more dispensable that those providing functions host genomes are not capable of (Fellous and Salvaudon 2009). Besides, it could be argued that the fitness effects of alternative plastic strategies may depend on the environmental context more than general improvement of resource availability (Chevin et al 2010). Therefore, symbionts that improve general performance of the host through greater resource availability may be more likely to be fixed among host individuals and populations than those that affect plasticity. By contrast, hosts may dynamically acquire and lose symbionts which effects on fitness depend on the environment, paving the way for the evolution of facultative symbiosis. Along these lines, recent modelling of host-symbiont dynamics revealed that whether symbionts affect adult survival or reproduction determines transmission mode evolution (Brown and Akçay 2019). Our experimental study only considered one trade-off between two developmental traits, possibly overlooking effects on other fitness components. Future analyses should increase in dimensionality and consider a greater number of fitness components. Similarly, a precise description of the slopes and shapes of considered trade-offs will be necessary to discriminate simultaneous effects of symbionts on plasticity and resource acquisition. We are now pursuing further investigation to determine how and when bacterial and yeast symbionts affect host developmental plasticity and resource availability in *Drosophila* flies.

### Context-dependent effects of bacteria enable symbiont-mediated adaptation

A consequence of *Drosophila* bacterial symbionts having different effects in different environments is the possibility they contribute to the fine-tuning of host phenotype to local conditions (Margulis and Fester 1991; Moran 2007; Sudakaran et al. 2017). The phenomenon is well established in vertically transmitted symbionts of insects that protect their hosts from parasites. For example, populations of aphids exposed to parasitoids harbor protective *Hamiltonella* symbionts at greater frequency than parasitoid-free populations (Oliver et al. 2005). Similarly, in the fly *Drosophila neotestacea*, the spread of the bacterium *Spiroplasma* allowed hosts to evolve greater resistance to parasitic nematodes (Jaenike et al. 2010). Vertically transmitted bacterial symbionts of *Paramecium* ciliates can also improve host resistance to stressful conditions (Hori and Fujishima 2003). Whether bacteria act as parasites or mutualists depends partly on the genetic ability of the host to deal with stress in absence of the symbiont (Duncan et al. 2010). However, the evolutionary role of symbionts that may be acquired from the environment is less clear, in part because the mechanisms favoring the association of hosts with locally beneficial symbionts are not as straightforward as for vertical transmission (Ebert 2013). Nonetheless, several lines of evidence suggest environmentally acquired microbial symbionts may contribute to local adaptation in *Drosophila*-microbe symbiosis. First, symbionts can be transmitted across metamorphosis (i.e. transstadial transmission from the larval to the adult stage) and pseudo-vertically during oviposition (i.e. from mothers to offspring) (Bakula 1969; Starmer et al. 1988; Spencer 1992; Ridley et al. 2012; Wong et al. 2015; Téfit et al. 2018). Second, host immune system participates in the destruction of harmful gut bacteria and the retention of beneficial ones (Lee et al. 2017; Lee et al. 2018). Third, *Drosophila* larvae may preferentially associate with beneficial yeast species ensuring they engage in symbiosis with locally adequate nutritional symbionts (Fogleman et al. 1981; Fogleman et al. 1982). In addition to host genetic and preferential association with beneficial microbes, *Drosophila* adaptation to local conditions thanks to microorganisms further necessitates symbionts have different effects in different environments. Our results show bacteria isolated from a fly population have different effects on host phenotype depending on the substrate larvae were reared in (Figures 2, 3, 4 and 5). It is therefore possible that, in the field, locally beneficial extracellular bacterial symbionts contribute to *Drosophila* local adaptation through variations in symbiont community composition.

## Conclusion

In this study, we found that associations between laboratory *Drosophila* flies and their microbial symbionts result in different effects on host phenotype when the symbiosis is investigated under laboratory conditions or under conditions more comparable to natural ones. The context-dependency of bacterial effects and the underlying mechanisms we unveiled (i.e. bacterial ecology and bacterial effects on host plasticity) shed light on the role of microorganisms in the evolution of their hosts. While the universality of our results is limited by the use of laboratory insects and bacteria, they point out that in order to understand the ecology and evolution of symbiotic interactions in the wild it is necessary to use ecologically realistic conditions, which is attainable in the *Drosophila* system.

## Acknowledgements

We warmly thank L. Benoit and P. Gautier for methodological help and S. Bourg, M.P. Chapuis, S. Charlat, J. Collet, D. Duneau, O. Duron, R. Gallet, P. Gautier, N. Kremer, F. Leulier, N. Rode and F. Vanlerberghe for useful comments on an earlier version of this work. Version 3 of this preprint has been peer-reviewed and recommended by Peer Community In Evolutionary Biology (https://doi.org/10.24072/pci.evolbiol.100085).

## Supplementary Materials

### Supplementary Material 1. Laboratory recipes

**Table S1A:** laboratory medium recipe.

| Component | *Amount for 1.5L* |
| --- | --- |
| Reverse osmosis water | 1200 ml |
| Banana | 280 g |
| Sugar | 74 g |
| Dead yeast | 74 g |
| Alcohol | 30 ml |
| Agar | 12 g |
| Nipagin | 6 g |

**Table S1B:** lysogeny broth (LB) recipes.

| Component | *Quantity / Volume for* |  |  |  |
| --- | --- | --- | --- | --- |
|  | *Liquid LB* | *Agar LB* | *Anti-bacteria*<br>*Agar LB* | *Anti-yeast*<br>*Agar LB* |
| Reverse osmosis water | 1000 ml | 1000 ml | 1000 ml | 1000 ml |
| Proteose peptone n°3 (Conda) | 10 g | 10 g | 10 g | 10 g |
| Yeast extract (Merck) | 5 g | 5 g | 5 g | 5 g |
| NaCl (Carlo Erba) | 5 g | 5 g | 5 g | 5 g |
| European Bacteriological Agar (Conda) |  | 15 g | 15 g | 15 g |
| Ampicillin (Sigma) (pure) |  |  | 100 mg |  |
| Chloramphenicol (Sigma) (100 mg/ml in ethanol) |  |  | 10 mg |  |
| Cycloheximide (Sigma) (100 mg/ml in DMSO) |  |  |  | 1 mg |

### Supplementary Material 2. Bacterial strains isolated from Oregon-R *Drosophila melanogaster* and used in the experiment

**Figure S2:**
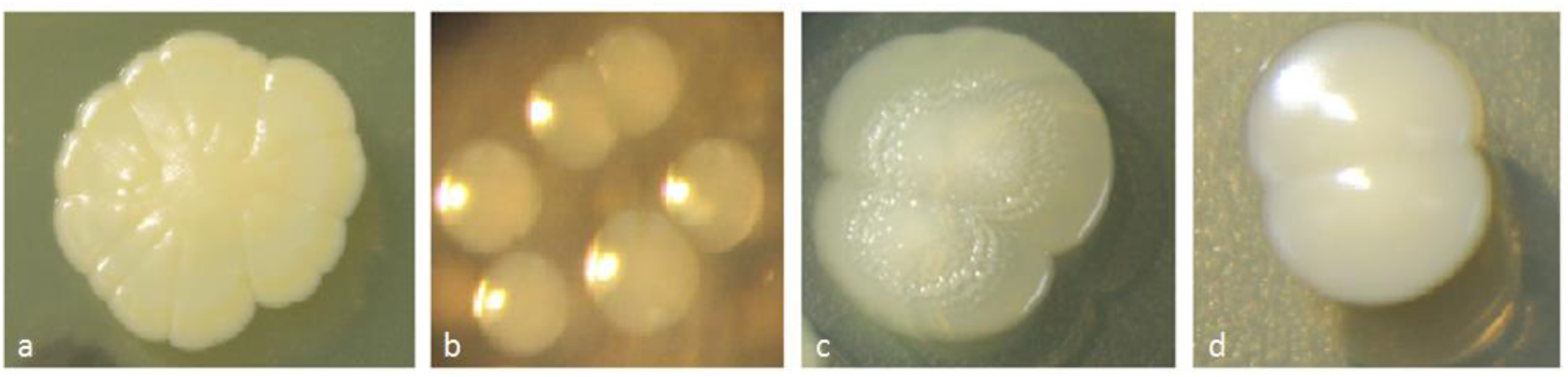
bacterial strains isolated from Oregon-R *Drosophila melanogaster* and used in the experiment. (a) *Staphylococcus* sp.; (b) *Enterococcus* sp.; (c) Enterobacteriaceae; (d) Actinobacteria.

### Supplementary Material 3. Live yeast as a prerequisite to *D. melanogaster* larvae survival on pristine grape berry

**Figure S3:**
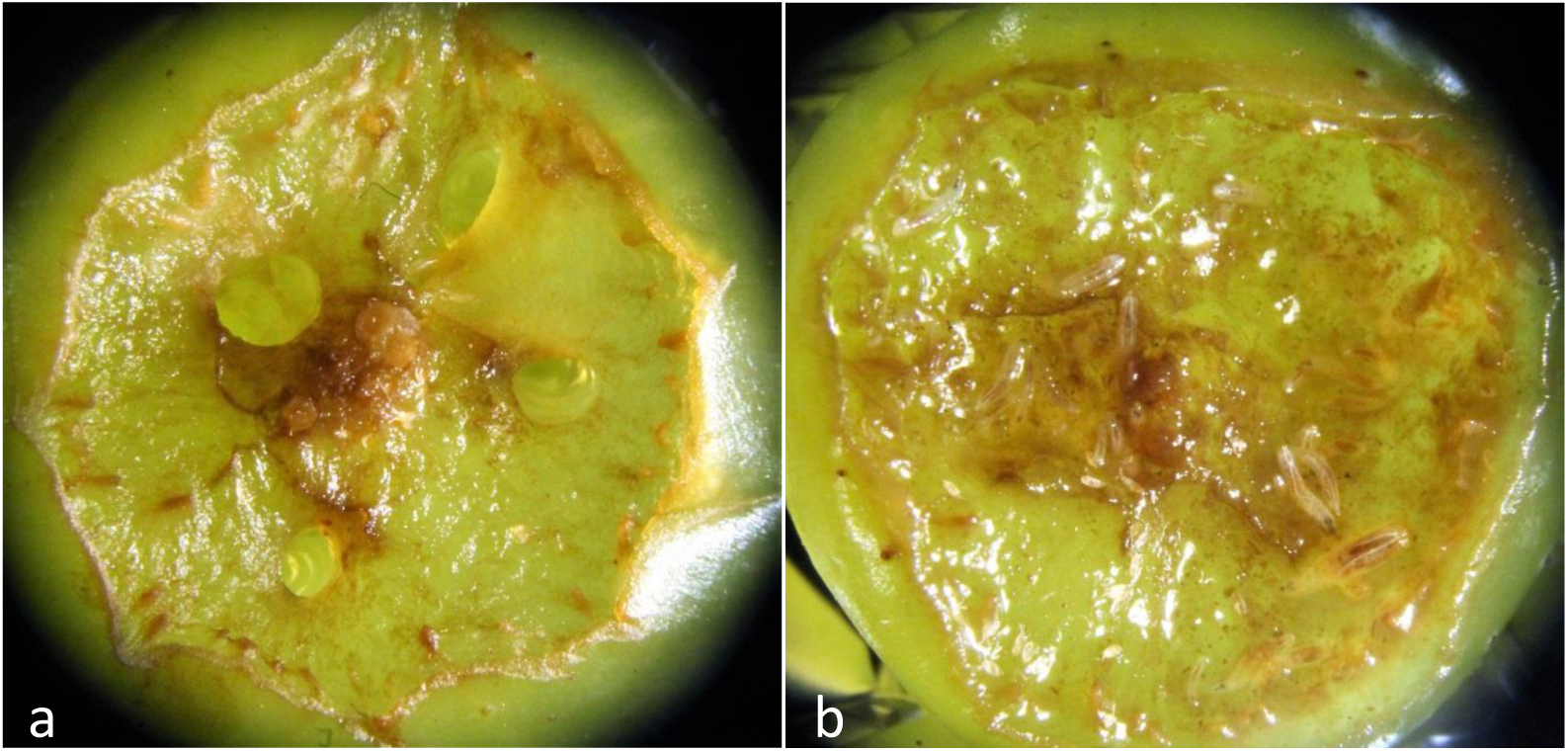
live yeast is necessary for the survival of *D. melanogaster* larvae on pristine grape berry. Prior to the experiment, we investigated survival of *D. melanogaster* larvae on fresh grape berries. Twenty bacteria-free *D. melanogaster* eggs were deposited next to an artificial wound with or without the bacterial isolates and *Saccharomyces cerevisiae*. In absence of yeast, larvae died quickly after hatching, with or without bacteria (Figure S1a). When live yeast was added to the system, numerous larvae developed up to the 3^thrd^ instar (Figure S1b), when we stopped monitoring.

### Supplementary Material 4. Experimental design for the grape berry environment

**Figure S4:**
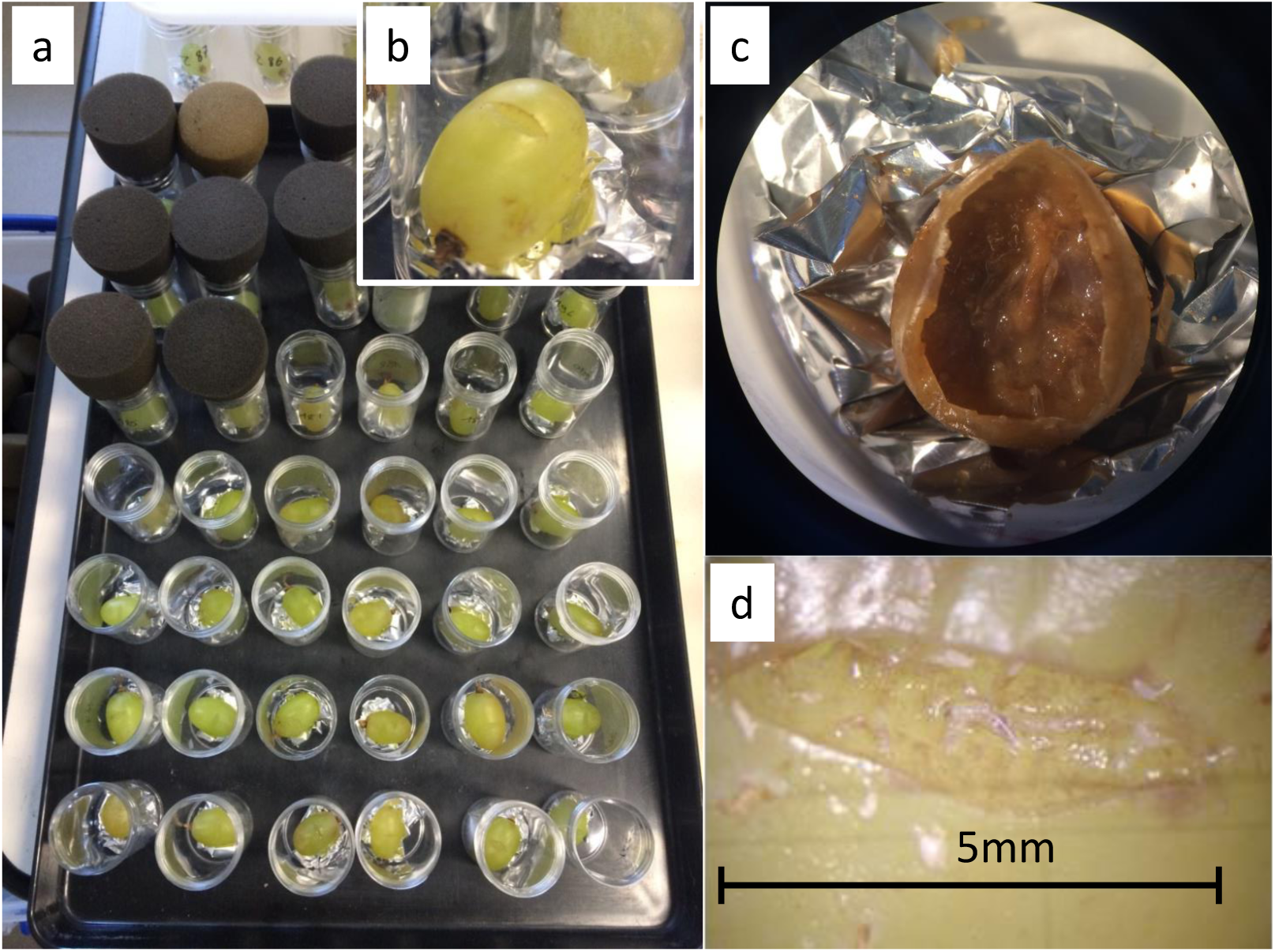
experimental design for the grape berry environment. (a) Experimental block for grape berry treatments, (b) Experimental unit with grape berry, (c) Decaying grape berry with live yeast, bacteria and larvae, (d) Egg cases visible near berry incision and active larvae in fruit flesh.

### Supplementary Material 5. Bacterial physiological profiles

#### Text S5

Eco Microplates (Biolog) were used to have an overview of the metabolic ‘fingerprint’ of the Enterobacteriaceae, the Actinobacteria isolate and the Actinobacteria variant. A fixed number of fresh bacteria cells suspended in sterile PBS were inoculated in well with one of 31 different carbon sources. Each combination Bacterial isolate*Carbon source was replicated three times. The plates were incubated at 25 °C and the absorbance at 595 nm was measured with a Multiskan GO spectrometer (Thermo Scientific) after 48 h and 120 h. A tetrazolium dye included with each carbon source entrained the production of red color when bacterial respiration occurred, i.e. when the carbon source was used. Variations of red color among carbon sources allowed establishing a physiological profile of each bacterial isolate.

**Figure S5A:**
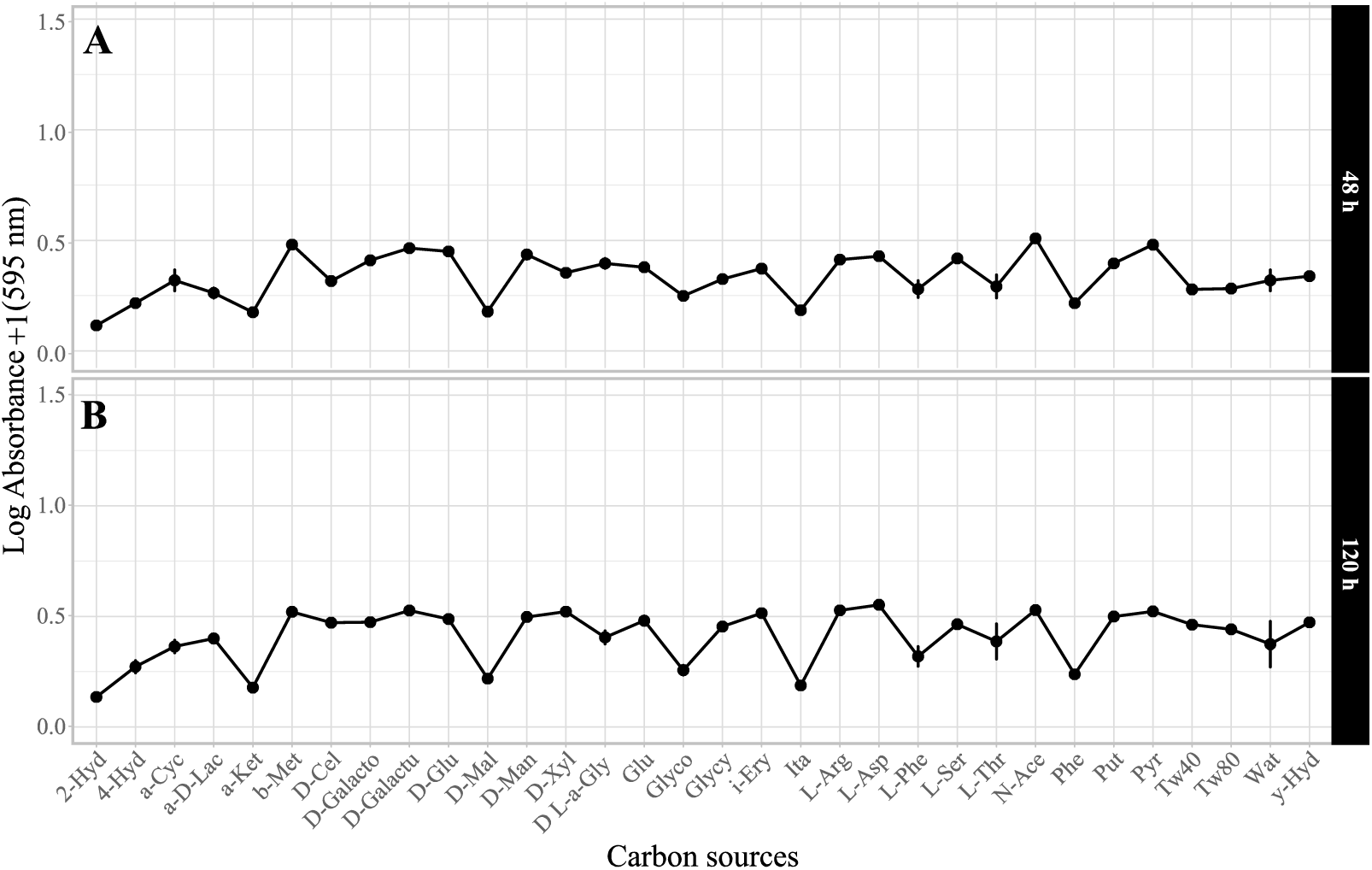
physiological profile of the Enterobacteriaceae isolate after (A) 48 h- and (B) 120 h-long exposure to different carbon sources. Symbols indicate means; error bars indicate standard errors around the mean. X-axis labels correspond to abbreviations of tested carbon sources, with 2-Hyd for 2-Hydroxy Benzoic Acid; 4-Hyd for 4-Hydroxy Benzoic Acid; a-Cyc for α-Cyclodextrin; a-D-Lac for α-D-Lactose; a-Ket for α-Ketobutyric Acid; b-Met for β-Methyl-D-Glucoside; D-Cel for D-Cellobiose; D-Galacto for D-Galactonic Acid γ-Lactone; D-Galactu for D-Galacturonic Acid; D-Glu for D-Glucosaminic Acid; D-Mal for D-Malic Acid; D-Man for D-Mannitol; D-Xyl for D-Xylose; D L-a-Gly for D,L-α-Glycerol Phosphate; Glu for Glucose-1-Phosphate; Glyco for Glycogen; Glycy for Glycyl-L-Glutamic Acid; i-Ery for i-Erythritol; Ita for Itaconic Acid; L-Arg for L-Arginine; L-Asp for L-Asparagine; L-Phe for L-Phenylalanine; L-Ser for L-Serine; L-Thr for L-Threonine; N-Ace for N-Acetyl-D-Glucosamine; Phe for Phenylethylamine; Put for Putrescine; Pyr for Pyruvic Acid Methyl Ester; Tw40 for Tween 40, Tw80 for Tween 80, Wat for Water and y-Hyd for γ-Hydroxybutyric Acid.

**Figure S5B:**
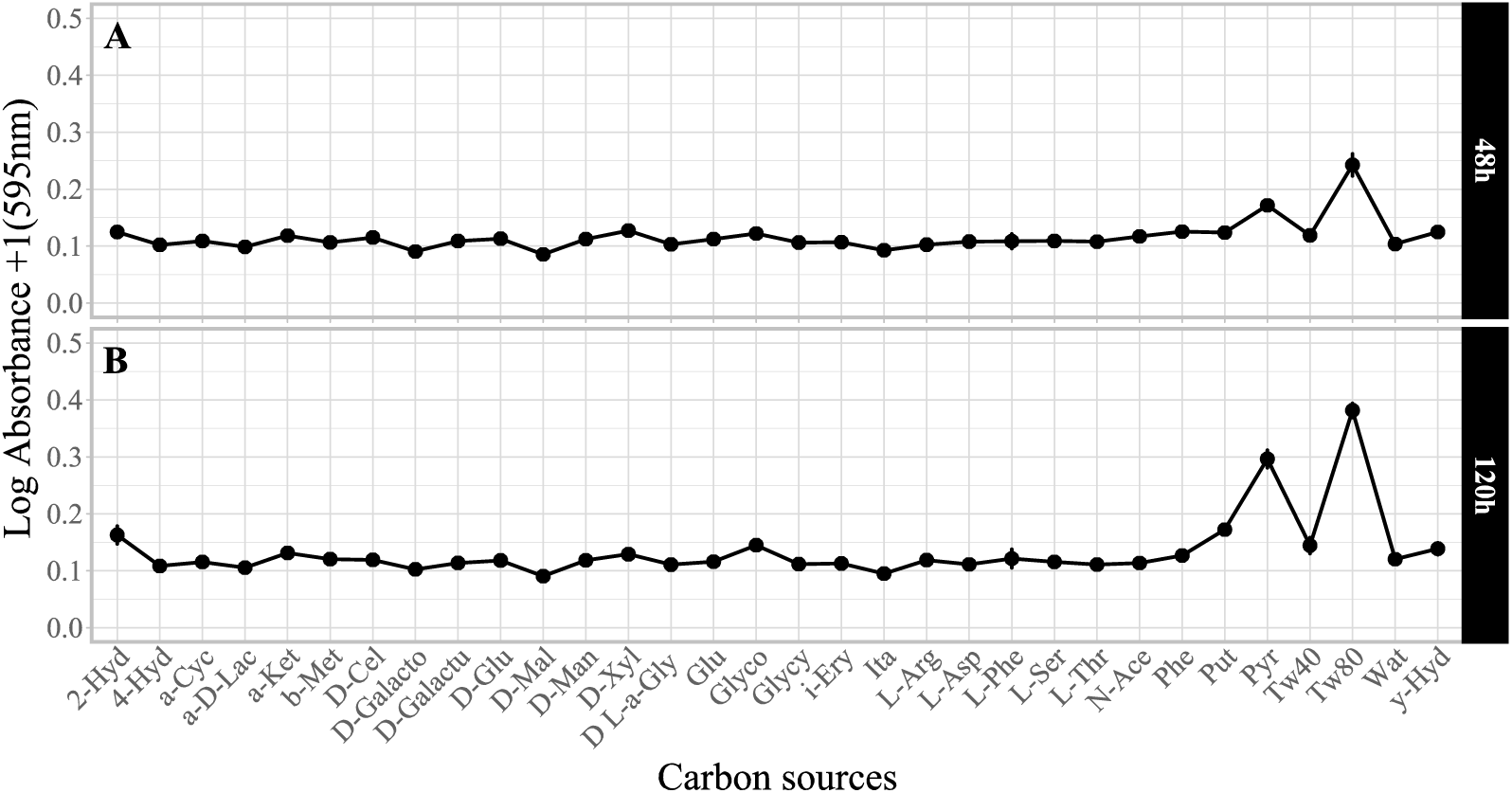
physiological profile of the Actinobacteria isolate after (A) 48 h- and (B) 120 h-long exposure to different carbon sources. Symbols indicate means; error bars indicate standard errors around the mean. X-axis labels correspond to abbreviations of tested carbon sources, with 2-Hyd for 2-Hydroxy Benzoic Acid; 4-Hyd for 4-Hydroxy Benzoic Acid; a-Cyc for α-Cyclodextrin; a-D-Lac for α-D-Lactose; a-Ket for α-Ketobutyric Acid; b-Met for β-Methyl-D-Glucoside; D-Cel for D-Cellobiose; D-Galacto for D-Galactonic Acid γ-Lactone; D-Galactu for D-Galacturonic Acid; D-Glu for D-Glucosaminic Acid; D-Mal for D-Malic Acid; D-Man for D-Mannitol; D-Xyl for D-Xylose; D L-a-Gly for D,L-α-Glycerol Phosphate; Glu for Glucose-1-Phosphate; Glyco for Glycogen; Glycy for Glycyl-L-Glutamic Acid; i-Ery for i-Erythritol; Ita for Itaconic Acid; L-Arg for L-Arginine; L-Asp for L-Asparagine; L-Phe for L-Phenylalanine; L-Ser for L-Serine; L-Thr for L-Threonine; N-Ace for N-Acetyl-D-Glucosamine; Phe for Phenylethylamine; Put for Putrescine; Pyr for Pyruvic Acid Methyl Ester; Tw40 for Tween 40, Tw80 for Tween 80, Wat for Water and y-Hyd for γ-Hydroxybutyric Acid.

### Supplementary Material 6. Joint analysis of bacterial effects on adult age and size for each sex

**Figure S6:**
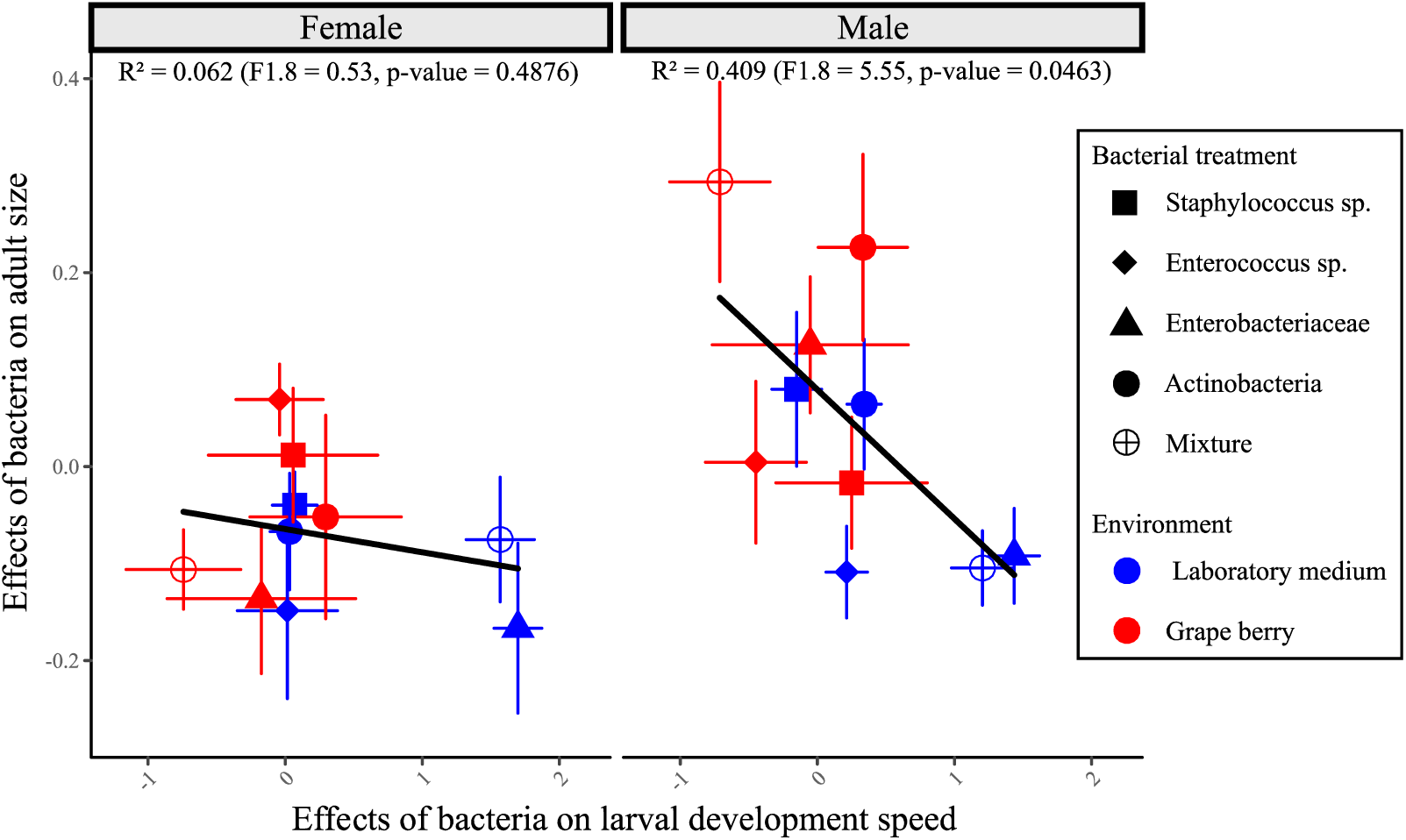
relationship between bacterial effects on age of emerging adults and bacterial effects on adult size, in females and males. As the linear regressions were not significantly different (Interaction Sex*Speed: F_1.16_ = 2.93, p = 0.11), data was pooled for the analysis reported in the main text (Figure 5). Symbols indicate the phenotype mean of each combination of bacterium and environment. Error bars mark the SE of the mean for both axes.

**Table S6:** Multivariate Analysis of Variance of the joint effect of the bacteria on ‘Age at emergence’ and ‘Adult size’ for each sex. As in Figure S6, general effects of the environments were removed by subtracting trait values of controls (i.e. without bacterial addition) in each environment before carrying out the analysis.

| *Factor* | *F* | *d.f.* | *p* |
| --- | --- | --- | --- |
| Environment | 8.72 | 1.78 | 0.0042 |
| Bacterial treatment | 0.73 | 4.78 | 0.57 |
| Environment*Bacterial treatment | 4.48 | 4.78 | 0.0026 |

**(b) *D. melanogaster* male**
| *Factor* | *F* | *d.f.* | *p* |
| --- | --- | --- | --- |
| Environment | 5.5 | 1.80 | 0.022 |
| Bacterial treatment | 1.56 | 4.80 | 0.19 |
| Environment*Bacterial treatment | 2.12 | 4.80 | 0.086 |

### Supplementary Material 7. Laboratory medium inoculated with the Enterobacteriaceae

**Figure S7:**
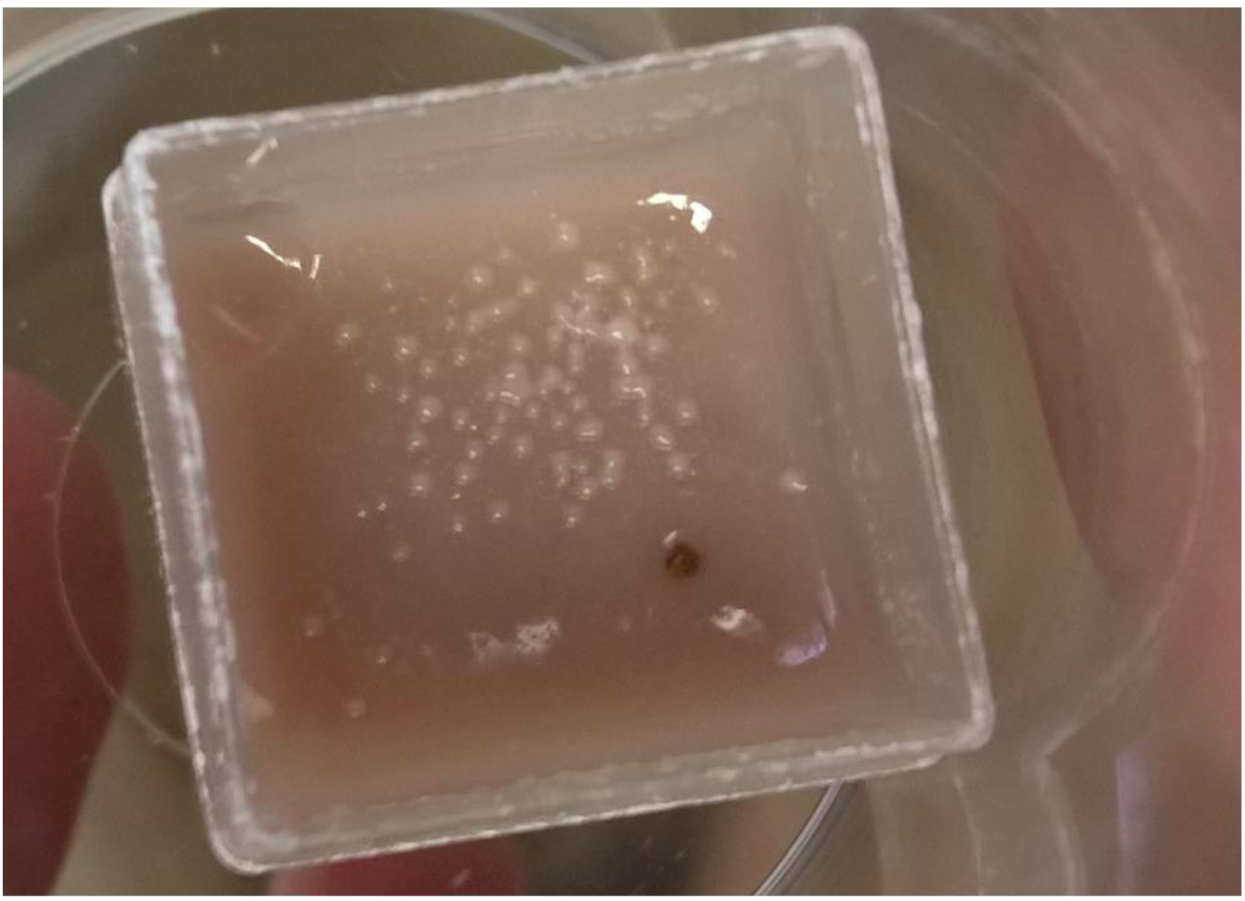
bacterial growth at the surface of laboratory medium five days after Enterobacteriaceae inoculation. This picture was taken in absence of larvae, but similar growth could be observed in their presence.

